# Deoxynivalenol induced inflammation and increased the adherence of entero-invasive *Escherichia coli* to intestinal epithelial cells via modulation of mucin and pro-inflammatory cytokine production

**DOI:** 10.1101/2024.03.11.584405

**Authors:** Murphy LY Wan, Vanessa Co Anna, Paul C Turner, Shah P Nagendra, Hani El-Nezami

## Abstract

Deoxynivalenol (DON) is a mycotoxin that commonly occurs in crops. It was hypothesized that DON could trigger intestinal inflammation and increase the susceptibility of intestinal epithelial cells (IECs) to pathogen infection. Accordingly, the aim of this study was to investigate the effects of DON on intestinal susceptibility to pathogen infection. Semiconfluent Caco-2 cells were exposed to DON followed by acute entero-invasive *Escherichia coli* (EIEC) infection. The effects of DON and EIEC contamination on mucin, cytokines and related signal transduction pathways were examined as part of the local immune system. Caco-2 cells were able to generate a rapid immune response against DON with or without EIEC post-challenge. An increase in EIEC attachment to DON-exposed cells was observed, probably in part, mediated by modulation of secretory MUC5AC mucins and membrane bound MUC4 and MUC17 mucins. Cells were also able to express and produce important mediators of inflammation, such as cytokines as a result of activation of toll-like receptors signalling cascades, modulation of nuclear factor κ-light chain-enhancer of activated B cells (NK-κB) and/or mitogen-activated protein kinase (MAPK) pathways. These data indicate that DON may exert immunomodulatory effects on intestinal epithelial cells, which might thereby modify the susceptibility to bacterial infection.

## 1. Introduction

The mycotoxin deoxynivalenol (DON) is secondary metabolite of *Fusarium* fungi that frequently contaminate both human food and animal feed. Human ingestion of DON contaminated grains can lead to different diseases, including nausea, vomiting, diarrhoea, abdominal pain, headache, dizziness, and fever (Sobrova et al. 2010).

DON has been known to be immunotoxic affecting cells of the immune system and the gastrointestinal tract. DON induces “ribotoxic stress response”, which results in the elevation of many pro-inflammatory genes such as cyclooxygenase-2 (COX-2) and inducible nitric oxide synthase (iNOS) as well as cytokines such as interleukin (IL)-6, IL-1β and tumour necrosis factor (TNF)-α (Moon et al. 2002, Moon et al. 2003, Pestka et al. 2004).

Foodborne pathogens can cause severe diseases including diarrhoea, which are of considerable public health concerns. The most common foodborne pathogens found to contaminate food include *Escherichia coli* (*E. coli*) spp., *Clostridium perfringens*, *Listeria spp.* and *Salmonella enterica* serotypes Enteritidis (Maciorowski et al. 2007). Among them, the most important foodborne pathogen is *E. coli*. Entero-invasive *E. coli* (EIEC) are intracellular pathogens that invade colonic epithelial cells by first penetrating the intestinal epithelium through microfold cells (M cells) to gain access to the submucosa, producing a rather localized infection with subsequent destruction of the underlying mucosa as a result of their active replication (Croxen et al. 2013).

The mucus lining the intestinal epithelium provides an important physicochemical barrier against ingested pathogens and toxins. It is composed of a mixture of mucins, which are heavily glycosylated with O-linked oligosaccharides and N-glycan chains linked to protein backbones. There are two different structural and functional classes of mucins: secreted gel-forming mucins (MUC2, MUC5AC, MUC5B, MUC6 and MUC19), and transmembrane (membrane-bound) mucins (MUC1, MUC3A, MUC3B, MUC4, MUC12, and MUC17). To initiate infection processes, bacteria need to find ways to ensure their survival and colonization by first adhering to the intestinal epithelium (Torres et al. 2003). Mucins can effectively impede the ability of bacteria and viruses to invade and colonize the cells, prevent their spread along the mucosal surfaces, and limit the amount of microbial-produced toxins reaching mucosal cells (Liévin-Le Moal et al. 2006).

Previous studies from our laboratory and others have shown that DON affected mucus production (Wan et al. 2014, Antonissen et al. 2015, Pinton et al. 2015) and also cytokine production (Van De Walle et al. 2008, Van De Walle et al. 2010, Wan et al. 2013). Bacterial pathogens such as *E. coli* could also trigger host inflammatory responses by enhancing mucin expression that protect bacterial intrusion to gut epithelium (Vieira et al. 2010, Xue et al. 2014), and eliciting cytokine production (Jung et al. 1995, Eaves-Pyles et al. 2008). *E. coli* can also be recognized by toll-like receptors (TLRs) (mainly TLR4), which transduce signals to the nucleus, activate the transcription factor nuclear factor κ-light chain-enhancer of activated B cells (NF-κB) and initiate inflammatory responses by producing inflammatory cytokines and chemokines (Underhill et al. 2002, Dąbek et al. 2010). The intestinal tract is the first barrier to ingested mycotoxins but also the first line of defence against intestinal infection. Ingestion of some mycotoxins increases the susceptibility to experimental or natural mucosal infections (Tai et al. 1988, Fukata et al. 1996, Stoev et al. 2000, Oswald et al. 2003, Vandenbroucke et al. 2011). So far, there is only one study reporting that DON rendered the intestinal epithelium more susceptible to *Salmonella Typhimurium* with a subsequent potentiation of the inflammatory responses in the gut (Vandenbroucke et al. 2011).

Accordingly, it was hypothesized that DON could also increase the susceptibility of intestinal epithelial cells (IECs) to an entero-invasive pathogen infection and trigger intestinal inflammation. With DON and EIEC infections being emerging issues with possible deleterious consequences for both animal and human health and with the gastrointestinal tract being the primary target, the aim of this study was to investigate the effects of low and relevant concentrations of DON on intestinal susceptibility to acute EIEC infection. Of the cell line models, the Caco-2 cell line, originally isolated from human colon adenocarcinoma, in spite of lack of mucus production, is among the most commonly used *in vitro* model of intestinal epithelium to study bacterial adherence and invasion (Resta-Lenert et al. 2003, Ganan et al. 2010, Resta-Lenert et al. 2011, Khodaii et al. 2017). Caco-2 cells were grown at semi-confluence stages (2 to 3 days of culture) and used in this study, as they were substantially more susceptible to infection than confluent cells.

## 2. Materials and Methods

### 2.1 Chemicals and reagents

Minimum Essential Medium (MEM), foetal bovine serum (FBS) and all other cell culture reagents were obtained from Gibco-Life Technology (Eggenstein, Germany). DON was obtained from Sigma Chemical Company (St. Louis, MO, USA) and was dissolved in dimethyl sulfoxide (DMSO) purchased from Sigma and stored at -20 °C before use. In addition, phosphate buffered saline (PBS), sodium biocarbonate, diethylpyrocarbonate (DEPC) and chloroform are purchased from Sigma. RNAisoPlus was purchased from Takara (Otsu, Japan). HiScript^TM^ RT SuperMix for qPCR and AceQ qPCR SYBR Green Master Mix were obtained from Vazyme Biotech Co. (Piscataway, NJ, USA).

### 2.2 Cell line and culture conditions

Caco-2 cells (passages 22-31) obtained from the ATCC (HTB-37) were maintained at 37 °C, 5% CO_2_, 90% relative humidity in MEM+20% FBS. Routinely, cells were sub-cultured once a week using trypsin-EDTA (0.25%, 0.53 mM) and seeded at a density of 2×10^6^ cells per 180 cm^2^ flask. All cells were screened for mycoplasma contamination with a MycoAlert mycoplasma detection kit (Lonza, Basel, Switzerland) prior to use.

### 2.3 Bacteria preparation

Entero-invasive *Escherichia coli* (EIEC O29:NM) was from Prof Wei Chen from the State Key Laboratory of Food Science and Technology, Jiangnan University. EIEC were incubated overnight at 37 °C in Luria-Bertani (LB) broth until the stationary phase was reached. Subcultures of the overnight cultures in fresh medium were grown to a phase of exponential growth. Cells were centrifuged at 4,000 rpm (Beckman Coulter, Fullerton, CA; GS-6R centrifuge) for 5 min, wash with PBS twice and suspended in MEM+20% FBS to desired concentrations before adding to the epithelial cell layers.

### 2.4 Cell viability by CCK-8 assay

A CCK-8 colorimetric assay was performed to assess cell viability/cytotoxicity in response to different concentrations of DON without or with EIEC post-treatment. The CCK assay is a colorimetric assay based on the reduction of a tetrazolium salt, WST-8 (4-[3-(2-methoxy-4-nitrophenyl)-2-[4-nitrophenyl]-2H-5-tetrazolio]-1,3-benzene disulfonate sodium salt), to a water-soluble formazan by cellular NADH or NADPH (Ishiyama et al. 1997). The assay was performed following manufacturer’s instruction (Dojindo, Kunamoto, Japan). In brief, cells were seeded at 2×10^4^ cells/well in 96-well culture plates and allowed to grow for 3 days. At 3 days, cells were treated with different concentrations of DON (0, 2, 4, 8, 16 µM) for 24 h. After 24 h incubation, Caco-2 cells were treated with medium containing exponentially grown EIEC bacteria at a multiplicity of infection (MOI) of 250:1 to the cells for 1 h. After that, cells were washed and incubated in medium with gentamicin (50 μg/ml) for another 1 h. At the end of the incubation, CCK-8 solution (10 μl) was added to each well, and the cells were incubated at 37 °C for 1 h. The colour intensity (absorbance) was determined using a microplate reader (model 550, BioRad) at 450 nm. Cell viability was expressed as the percentage of the mean value normalized to the control (untreated cells). For each treatment, the mean value was obtained from at least six wells.

### 2.5 Invasion and adhesion assay

The number of cells in each of the 24-well plate following DON treatment was first determined by trypan blue (Gibco) exclusion assay for calculation of MOI. Invasion assay were performed using the gentamicin protection assay as described previously (Boudeau et al. 1999). Briefly, Caoc-2 cells were seeded in 24-well plates (1×10^5^ cells) and cultured for 2 days. After 2 days, cells were treated with different concentrations of DON (0, 8, 16 µM) for 24 h. After 24 h incubation, cells were treated with medium containing exponentially grown EIEC at a MOI of 250:1 to the cells. After 1 h of incubation at 37°C, cells were washed and incubated in medium with gentamicin (50 μg/ml) (for cells infected with invasive bacteria or uninfected controls) for another 1 h at 37°C. The cells were then lysed with 0.1% Triton X-100 (Sigma) in deionized water. Samples were diluted and plated onto LB agar plates to determine the colony forming unit (CFU) recovered from the lysed cells. In control experiments, gentamicin has no significant effect on any of the parameters measured. Furthermore, no significant bacterial overgrowth was observed over the duration of the experiment under all conditions tested. The percentage of invading bacteria was expressed as CFU_(invading bacteria)_ of infected cells divided by CFU_(total bacteria)_, normalized to the number of cells in each of the 24-well plate following different concentrations of DON treatment.

To determine the total number of cell-associated bacteria corresponding to adherent and intracellular bacteria, cells were lysed after 1 h infection period, and the bacteria were quantified as described above. The number of adhering bacteria was determined by subtracting the number of invading bacteria from the total number of cell-associated bacteria. The percentage of adhering bacteria was expressed as CFU_(adhering bacteria)_ of infected cells divided by CFU_(total bacteria)_, normalized to the number of cells in each of the 24-well plate following different concentrations of DON treatment.

### 2.6 Quantification of MUC protein expression

Expression of MUC protein by Caco-2 cells was studied using biotinylated wheat germ agglutinin (WGA) (Vector Labs; Burlingame, CA, USA). Briefly, Caco-2 cells were seeded onto 96-well plates and treated or not with DON with or without EIEC infection as described above and were fixed with 4% paraformaldehyde (PFA; Sigma) during 30 min at room temperature. After fixation, cells were washed two times with 0.05% PBS-Tween 20 (PBS-Tw). Cells were blocked for 1 h at room temperature in PBS supplemented with 2% BSA and 0.1% Triton-×100. Biotinylated-conjugated WGA (1:10 000 dilution) was then added to the wells for 1 h at room temperature followed by incubation with avidin peroxidase (1:10 000 dilution) for another hour. Cells were washed two times with PBS-Tw and 3,3’,5,5’-tetramethylbenzidine peroxidase (TMB) substrate (BioLegend; San Diego, CA, USA) was added. Reaction was stopped with 2N sulphuric acid (Merck; Darmstadt, Germany) and the optical density was read at 490 nm using the Multiskan microplate spectrophotometer (ThermoFisher Scientific, Waltham, MA, USA). Relative MUC protein expression was normalized to the number of cells in the well following each treatment.

### 2.7 Quantitative polymerase chain reaction (qPCR) analysis

Cells were seeded and treated as above. After 1 h infection with EIEC, total RNA was extracted using RNAiso^TM^ Plus according to manufacturer’s instructions. The concentration of RNA was measured by using NanoDrop ND-1000 Spectrophotometer (Nano-Drop Technologies, Wilmington, DE) with purity ascertained by (A260/A280) of >1.8. RNA integrity was checked by running the RNA sample on ∼1% agarose gel. Total RNA (500 ng) from each sample was converted into cDNA using HiScript^TM^ RT SuperMix for qPCR according to the manufacturer’s instructions. qPCR was performed to quantify the products of interest, cytokines and chemokines (IL-1β, IL-6, IL-8, TNF-α, MCP-1), mucins (MUC1, MUC3A, MUC4, MUC5AC, MUC5B, MUC17), Toll-like receptors (TLR-1, -2, -4, -5, -6) and related signalling molecules (MyD88, NF-κB). Assessment of glyceraldehyde-3-phosphate dehydrogenase (GAPDH) levels was also performed which served as an internal control for RNA integrity and loading. Human specific primers were described in Table 1. For analyses on a StepOnePlus^TM^ Real-Time PCR system (Applied Biosystems, Foster City, CA, USA), 2 μl of 5X diluted cDNA was added to AceQ qPCR SYBR Green Master Mix, to obtain final primer concentrations of 500 nM/primer in final volume of 10 ul. The sample was centrifuged briefly and run on the PCR machine using the default fast program (45 cycles of 95°C for 3 s, 60 °C for 30 s). To ensure the reliability of qPCR data, the amplicons are kept short (<250 bp) (Nolan et al. 2006). All PCR reactions were performed in duplicate. Negative controls, consisting in PCR mix components without cDNA was used for all primers. The relative product levels were quantified using the 2^-△△CT^ method as described previously (Livak et al. 2001).

**Table 1.**
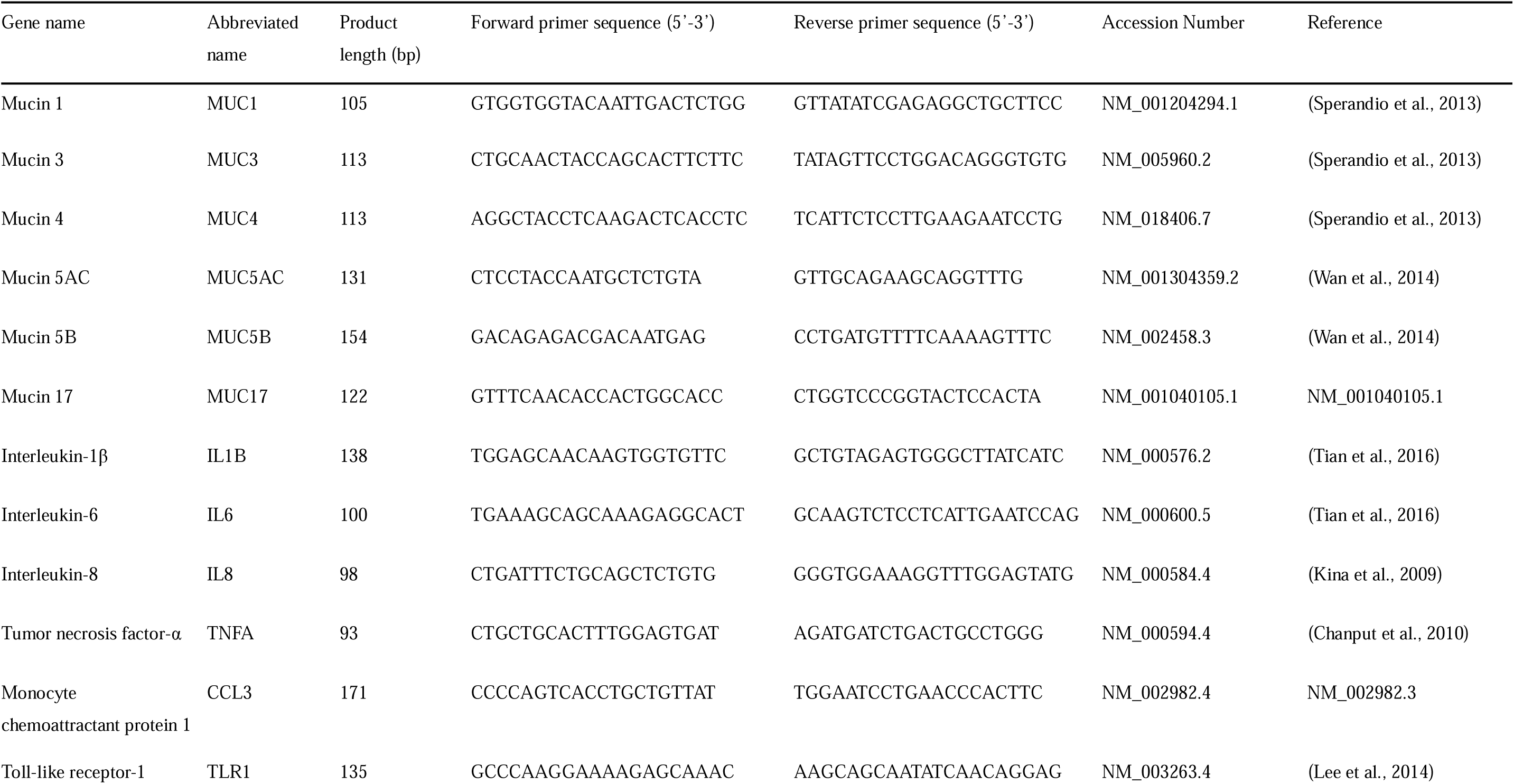
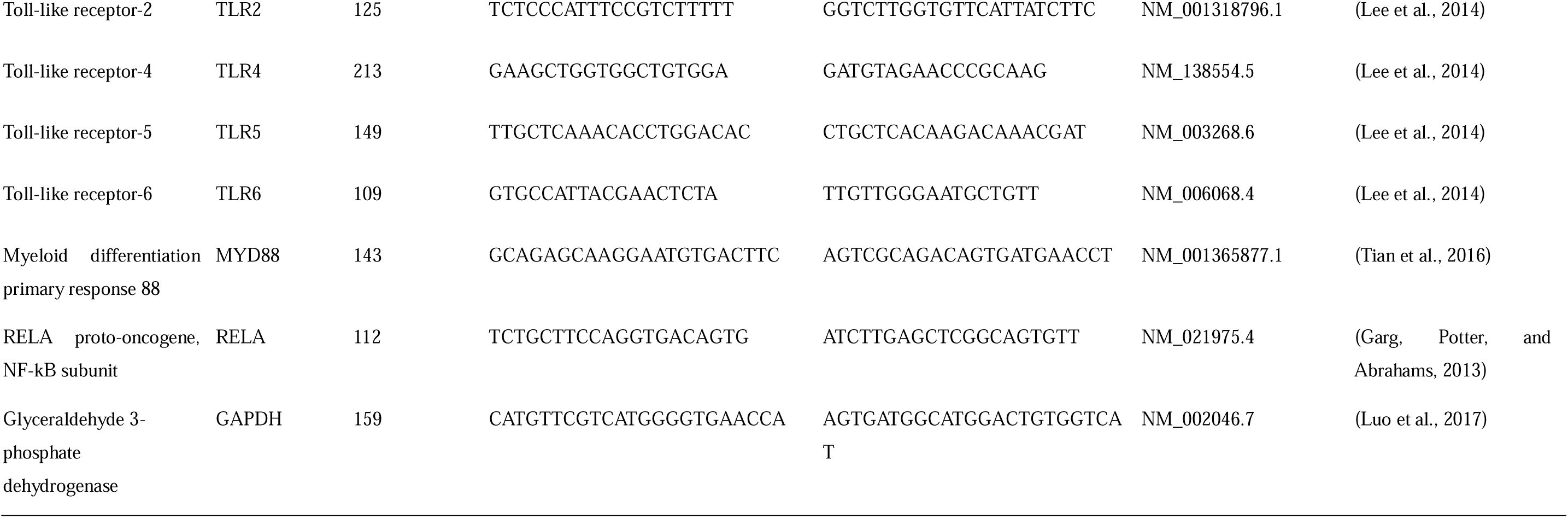
Human specific primer sequences for qPCR.

### 2.8 Protein extraction, SDS-PAGE and immunoblotting

The cells, undergoing the same treatment as described for qPCR, were washed with PBS and extracted for total proteins (whole cell extracts). In brief, 100 µl of RIPA lysis buffer, supplemented with protease inhibitor cocktail (Sigma) and phosphatase inhibitors (Cell Signaling, Beverly, MA, USA) was used to extract total proteins from the cells. Protein concentration was determined by the DC protein assay (BioRad).

Proteins (10-50 µg) were loaded onto 10% sodium dodecyl sulfate polyacrylamide gel (BioRad), separated by electrophoresis (SDS-PAGE), and then blotted onto a polyvinylidene difluoride (PVDF) membrane (Millipore, Darmstadt, Germany). The membrane was blocked with 5% BSA in Tris-buffered saline (TBS) containing 0.05% (v/v) Tween 20 (TBST) buffer. Proteins were probed by immunoblotting with diluted rabbit primary antibodies from Cell Signaling (1:1000) for NF-κB p65 (#8242), p44/42 MAPK (Erk1/2) (#4695), phospho-p44/42 MAPK (Erk1/2) (#4370), JNK2 (#9258), phospho-SAPK/JNK (#4668), p38 MAPK (#8690), phospho-p38 MAPK (#4511) and GAPDH (ab181602; Abcam), followed by horseradish peroxidase (HRP)-conjugated anti-rabbit IgG (#170-6515, BioRad) secondary antibodies. The blots were developed using Clarity Western ECL blotting kit (BioRad) and chemiluminescence was detected with a digital imaging system (ChemiDoc XRS+ system with image lab software, BioRad). Quantification was performed by Image Lab software version 6.0 (Bio-Rad) by densitometric analysis (Schneider et al. 2012).

### 2.9 Statistical analyses

All assays were expressed as mean ± standard error of mean (SEM) for the number of separate experiments indicated. Data analyses were performed using the GraphPad PRISM 9.0 software (Graphpad Software Inc., San Diego, CA). All data were first evaluated for normality with the Shapiro–Wilk and Levene’s variance homogeneity test. For parametric data, one-way analysis of variance (ANOVA) followed by Dunnet’s test against a control group; for non-parametric data, one-way ANOVA with the Kruskal-Wallis test, followed by the Dunn’s multiple comparisons test was used to identify significant differences against a control group. Data are significantly different at *p*<0.05, according to the post hoc ANOVA statistical analysis.

## 3. Results

### 3.1 DON and EIEC lowered cell viability

CCK assay was performed to investigate the effect of DON pre-treatment on cell viability and to determine the concentrations of DON used in subsequent experiments (Fig. 1A). DON resulted in a significant concentration-dependent reduction in cell viability at 4, 8 and 16 µM (*p*<0.05). DON at the concentrations of 8 and 16 µM were selected for the subsequent experimental assays. The combined effect of DON and EIEC on cell viability was also assessed. Addition of EIEC to cells pre-treated with 8 and 16 µM DON significantly reduced cell viability compared to control (PBS) (*p*<0.0001), but the reduction was not significantly different from cells treated with EIEC or DON alone (Fig. 1B).

**Fig. 1.**
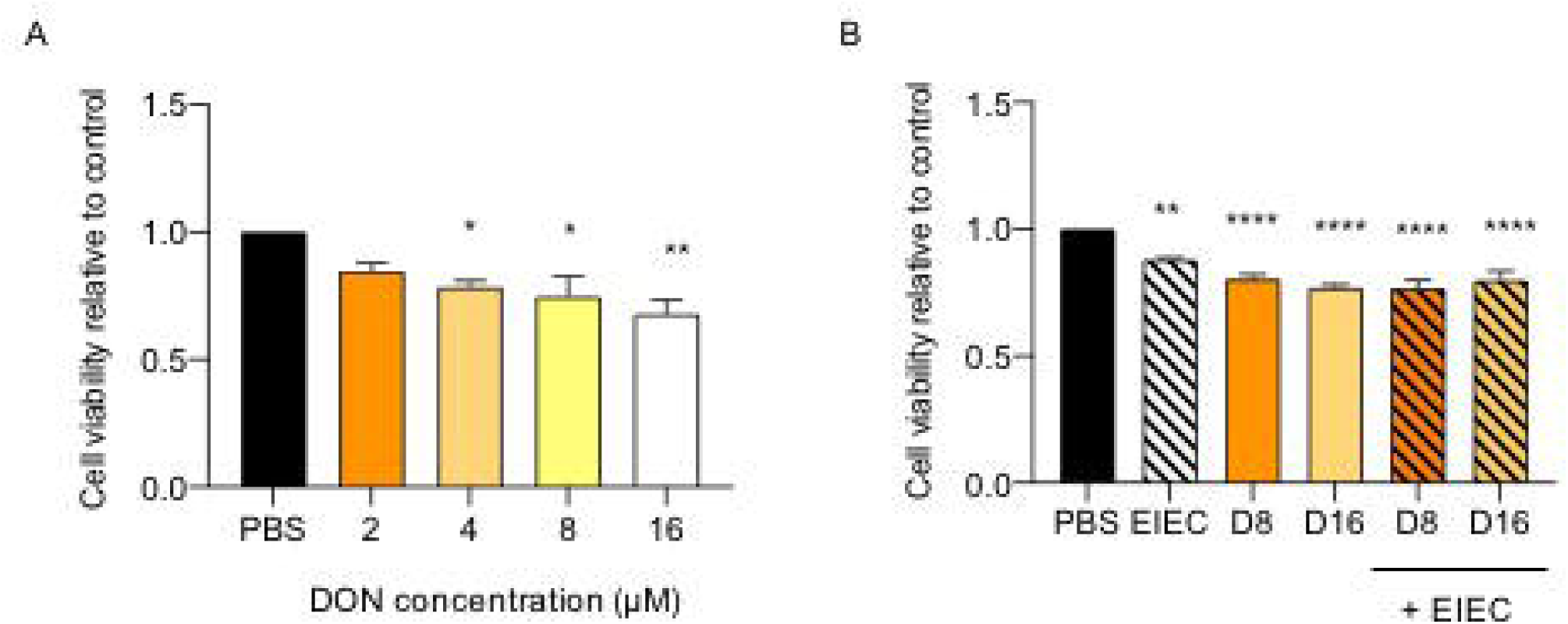
Effects of DON without or with EIEC post-infection on cell viability. (A) Preliminary screening of DON concentrations for subsequent experiments. Cell viability data of caco-2 cells treated with different concentrations of DON (0, 2, 4, 8 and 16 µM) for 24 hours. (B) Cell viability data of Caco-2 cells treated with DON (8 and 16 µM) for 24 hours without or with EIEC bacteria post-treatment at a multiplicity of infection (MOI) of 250:1 to the cells for 1 h. Control received appropriate carriers. Results were shown as mean ± SEM, which are from four independent experiments performed in six replicates. *, **, ***, *** *p*<0.05, 0.01, 0.001 and 0.0001 compared to PBS control. One-way ANOVA post Dunnet’s test.

### 3.2 DON increased EIEC adhesion but reduced invasion

Since adhesion and invasion are both important processes in bacterial pathogenesis, the effects of DON on EIEC adhesion and invasion were assessed by a modified bacterial adherence assay and gentamicin protection assay (Fig. 2). A preliminary bacterial invasion experiment was conducted in our laboratory by adding EIEC at MOI of 1:1, 2.5:1, 10:1, 25:1, 100:1, 250:1, 500:1, 1000:1 to the cells for 1 or 2 h. Results showed that EIEC at MOI of 250:1 for 1 h showed the highest invasion number to the cells from the colony counts (data not shown). Therefore, the MOI of 250:1 and 1 h of EIEC incubation were chosen for all the subsequent experiments. Pre-treatment with 8 and 16 µM DON caused a significant increase in EIEC adherence (*p*<0.05) (Fig. 2A). However, cells pre-treated with DON reduced EIEC invasion significantly at (*p*<0.05) (Fig. 2B).

**Fig. 2.**
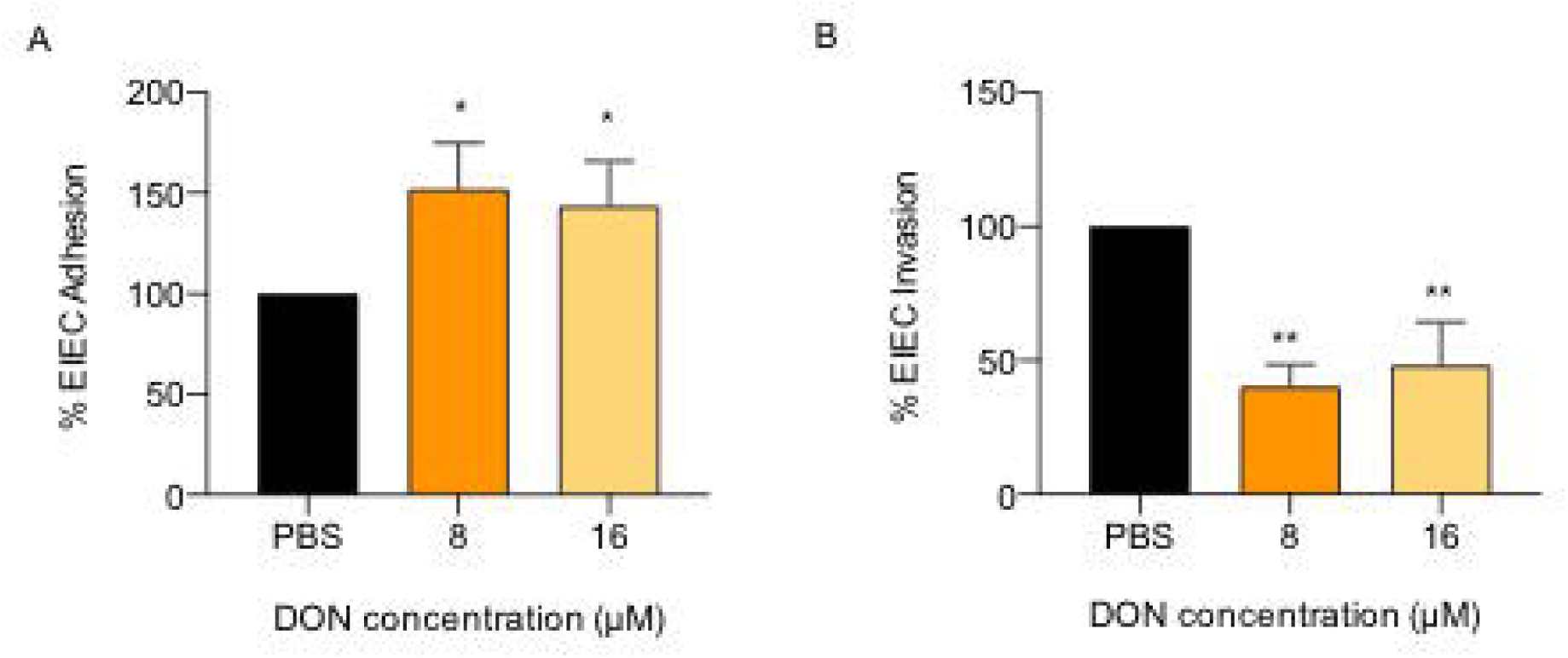
Effects of 24 h of DON incubation on Caco-2 cells without or with EIEC post-infection (1 h) on bacterial adhesion and invasion. The percentage of (A) adhering and (B) invaded bacteria were calculated as described in Materials and Methods. Results were shown as mean of ± SEM, which are from four separate experiments performed in duplicates. *, ** *p*<0.05 and 0.01 compared to PBS control. One-way ANOVA post Dunn’s test.

### 3.3 DON and EIEC contamination altered mucin gene expression and protein production

Mucin production is important in forming intestinal mucus, and acts as a physical barrier against bacterial infection. The effect of DON and EIEC exposure on mucin mRNA expression was determined by qPCR (Fig. 3) and protein production using the lectin wheat germ agglutinin (WGA) assay (Fig. 4).

**Fig. 3.**
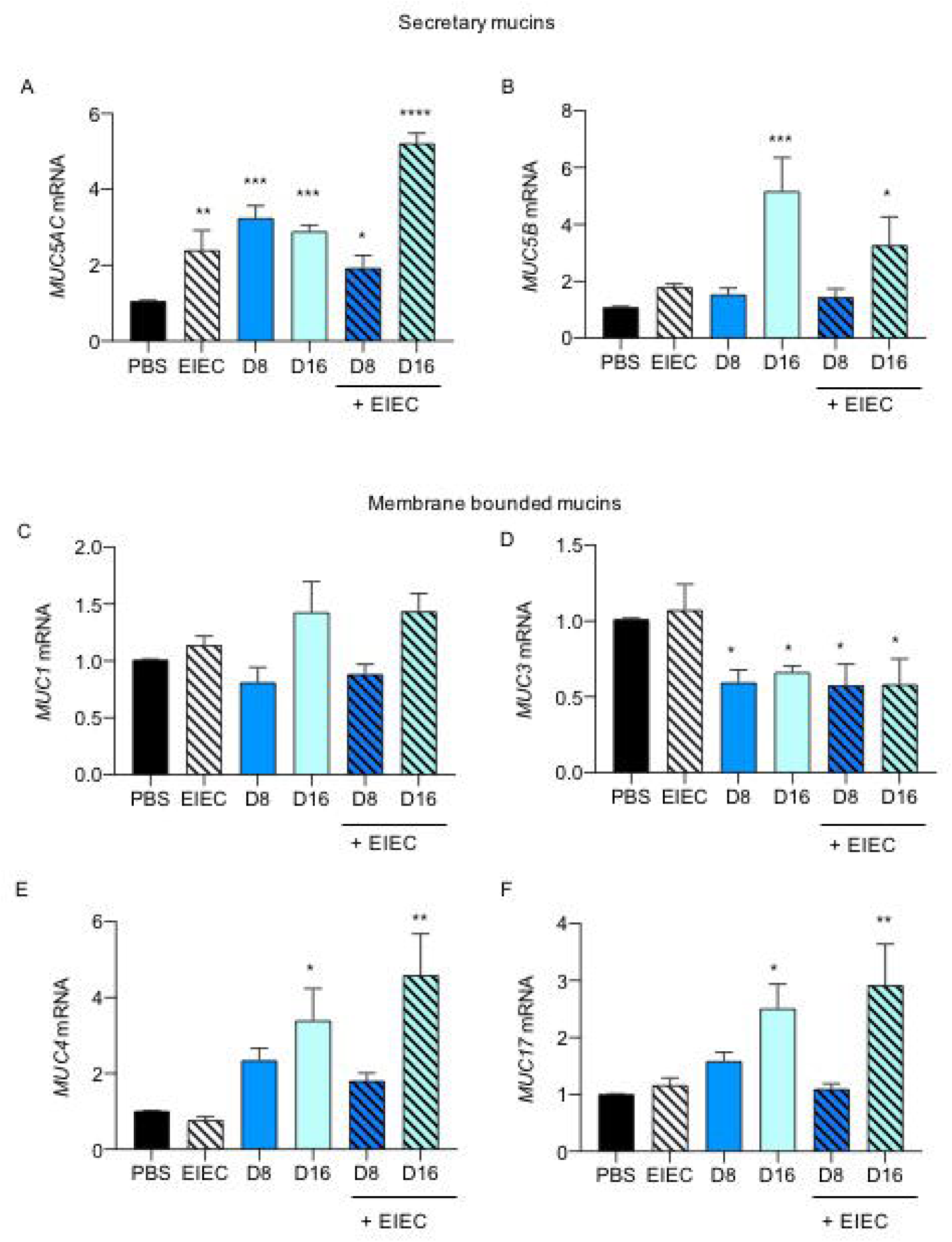
Effects of 24 h DON incubation without or with EIEC post-infection (1 h) on mucin (*MUC*) gene expression. (A-B) Secretory *MUC5AC* and *MUC5B*, and (C-F) membrane bound *MUC1, MUC3, MUC4* and *MUC17* mRNA expression was measured by qPCR, with *GAPDH* as the internal control. Results were shown as mean of ± SEM from five independent experiments. *, **, *** *p*<0.05, 0.01 and 0.001 compared to PBS control. One-way ANOVA post Dunnet’s test.

**Fig. 4.**
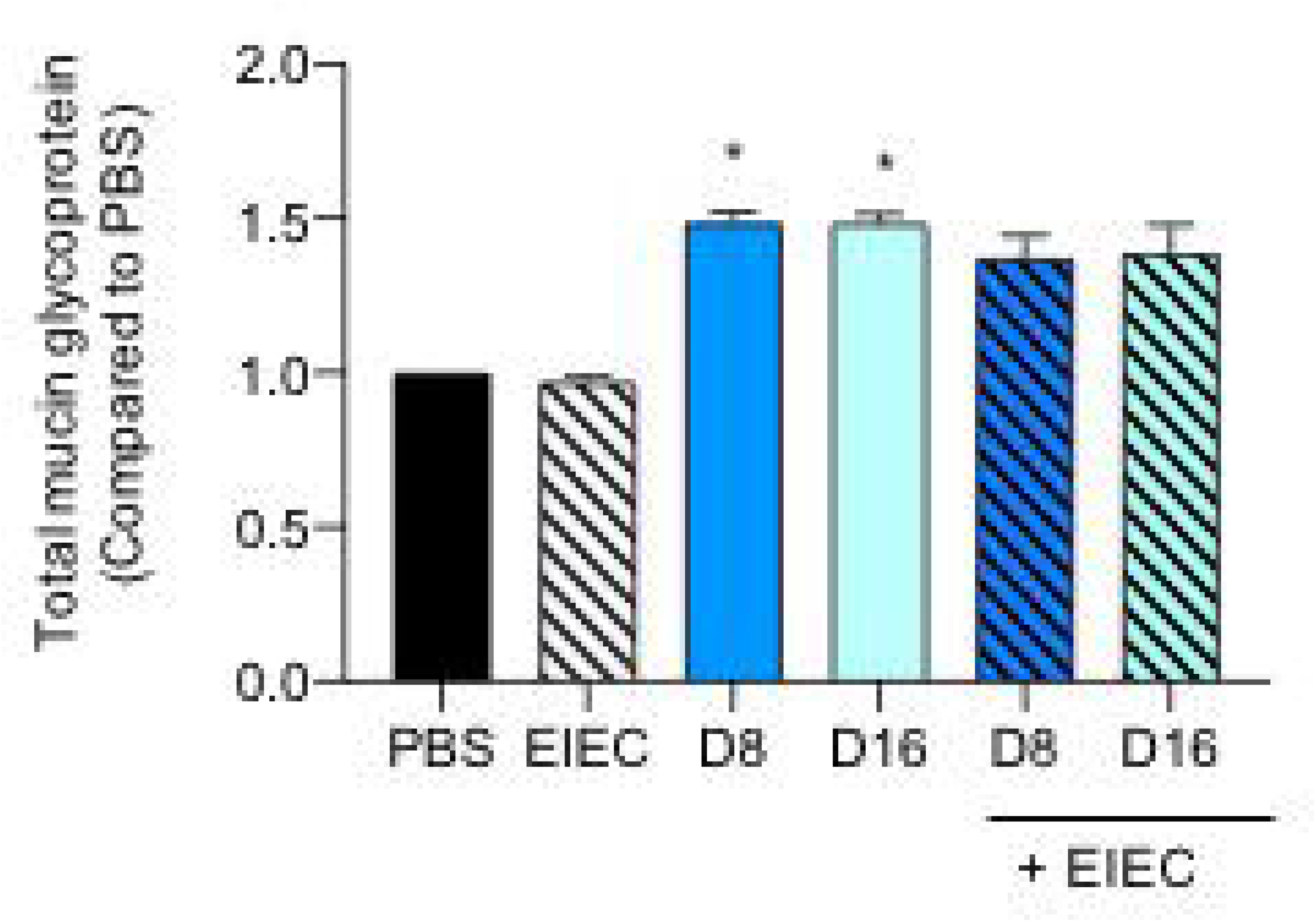
Effects of 24 h DON incubation without or with EIEC post-infection (1 h) on mucin (MUC) protein production as measured by WGA assay. Results were shown as mean of ± SEM from six independent experiments. * *p*<0.05 compared to PBS control. One-way ANOVA post Dunn’s test.

Significant up-regulation of *MUC5AC* mRNA was found when cells were treated with all the treatment groups (Fig. 3A). Significant up-regulation of *MUC5B* mRNA was also observed when cells were treated with 16 µM DON alone and with EIEC post-infection (Fig. 3B).

*MUC1*, *MUC4* and *MUC17* mRNA expression showed similar responses when treated with DON, where they experienced a gradual increase with increasing DON concentrations (Fig. 3C, E, F). For *MUC1*, in general, the mRNA expression level in cells with DON treatment alone remained unchanged when compared to PBS control. DON pre-treatment lowered *MUC3* mRNA expression level significantly (*p*<0.05). Addition of EIEC did not affect *MUC3* mRNA expressions (Fig. 3D). When Caco-2 was treated 16 µM DON, *MUC4* and *MUC17* mRNA expression increased significantly (*p*<0.05) compared to the PBS control. Addition of EIEC further increased *MUC4* and *MUC17* mRNA expression (*p*<0.01) (Fig. 3E-F).

To investigate the effects of DON and EIEC treatment on the mucus secretion, mucin-like glycoprotein secretion in cell lysates and those secreted into culture supernatants were measured and analysed. Exposure to 8 and 16 µM DON alone increased mucin protein production significantly (*p*<0.05). Addition of EIEC, however, did not cause any significant change to the amount of mucin protein produced compared to their pathogen-free counterparts (Fig. 4).

### 3.4 DON and EIEC modulated pro-inflammatory cytokine and chemokine gene expression

The effects of treating Caco-2 monoculture with DON and EIEC on pro-inflammatory cytokine and chemokine mRNA expression were examined by qPCR (Fig. 5). DON with and without EIEC post-treatment significantly downregulated *IL1B* gene expression (*p*<0.0001) (Fig. 5A). Pre-treatment with 16 µM DON upregulated *IL6* mRNA expression significantly (*p*<0.05); addition of EIEC further increased the *IL6* mRNA (*p*<0.001) (Fig. 5B). In the absence of DON, EIEC contamination alone significantly increased *IL8* and *TNFA* mRNA expressions (*p*<0.001 and 0.05, respectively) (Fig. 5C and D). Pre-treatment with all concentrations of DON lowered the *IL8* expression significantly (*p*<0.0001), but no significant change in *TNFA* was observed. Treatment with DON and EIEC both significantly down-regulated *CCL2* mRNA expression (*p*<0.001), though it appeared DON’s ability to modulate *CCL2* gene expression was greater than that of EIEC (*p*<0.0001) (Fig. 5E).

**Fig. 5.**
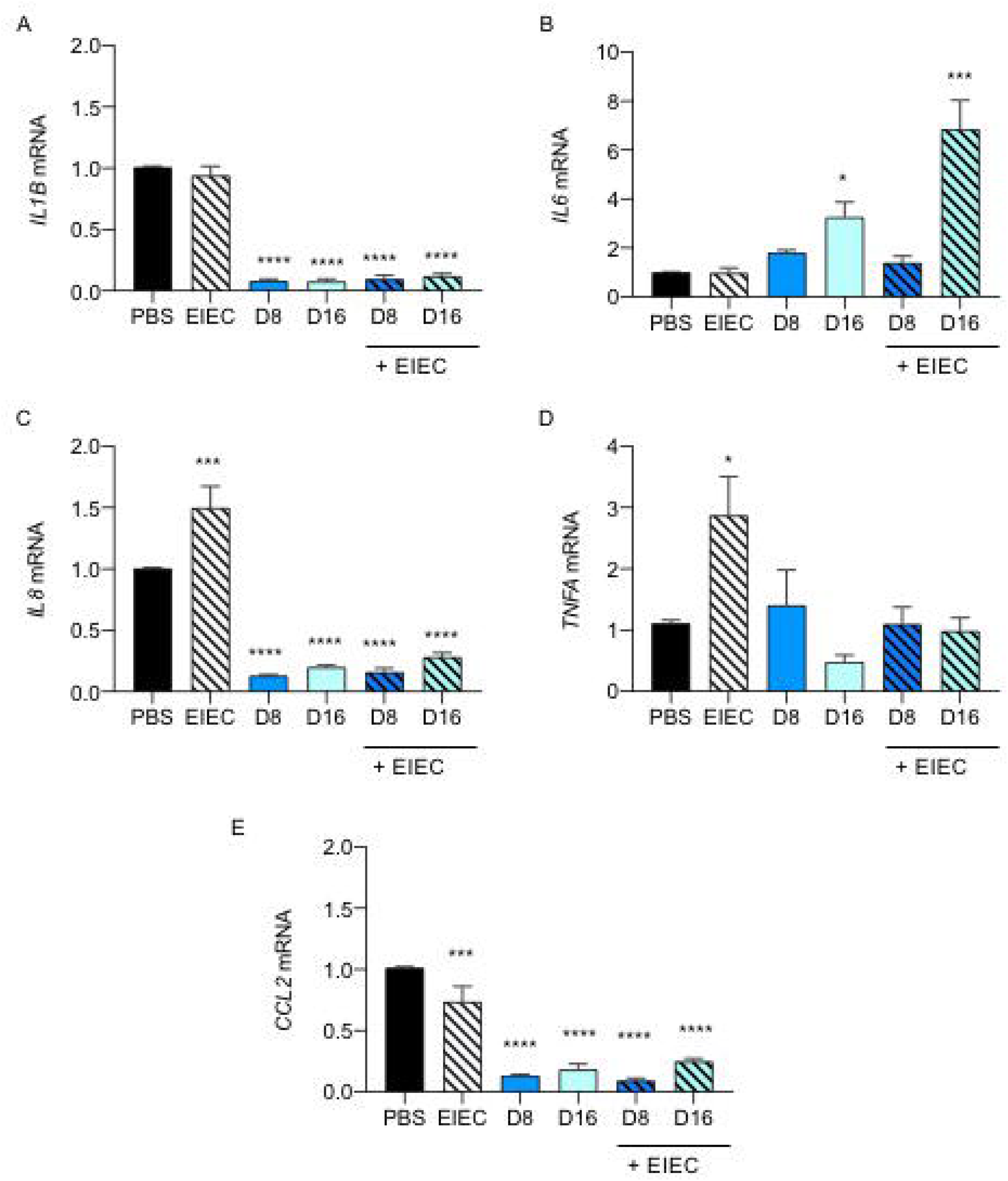
Effects of 24 h DON incubation without or with EIEC post-infection (1 h) on cytokine and chemokine gene expression. (A-E) *IL1B*, *IL6, IL8*, *TNFA* and *CCL2* mRNA expression was measured by qPCR, with *GAPDH* as the internal control. Results were shown as mean of ± SEM from five independent experiments. *, ***, **** *p*<0.05, 0.001 and 0.0001 compared to PBS control. One-way ANOVA post Dunn’s test.

### 3.5 DON and EIEC regulated TLRs and MyD88 gene expression

Pathogen associated molecular patterns (PAMPs) can be recognized by TLRs, which along with MyD88, can stimulate cytokines and chemokines production through NF-κB activation. TLRs and MyD88 gene expression in response to DON and EIEC were assessed by qPCR (Fig 6). DON showed modest upregulation of *TLR1* gene at all concentrations tested; addition of EIEC to 16 µM DON pre-treated cells, however, showed a remarkable increase in *TLR1* mRNA (*p*<0.001) (Fig. 6A). For *TLR5*, the gene expression was downregulated significantly by DON at all concentrations in the absence and presence of EIEC (*p*<0.0001) but no change was observed when cells were infected with EIEC alone (Fig. 6C). Due to the low expression of *TLR4* in Caco-2 cell line, its expression data for uninfected control and treatment groups were not shown. For *TLR2*, *TLR6* and *MYD88*, their gene expressions were not affected by the treatments significantly (Fig. 6B, D and E).

**Fig. 6.**
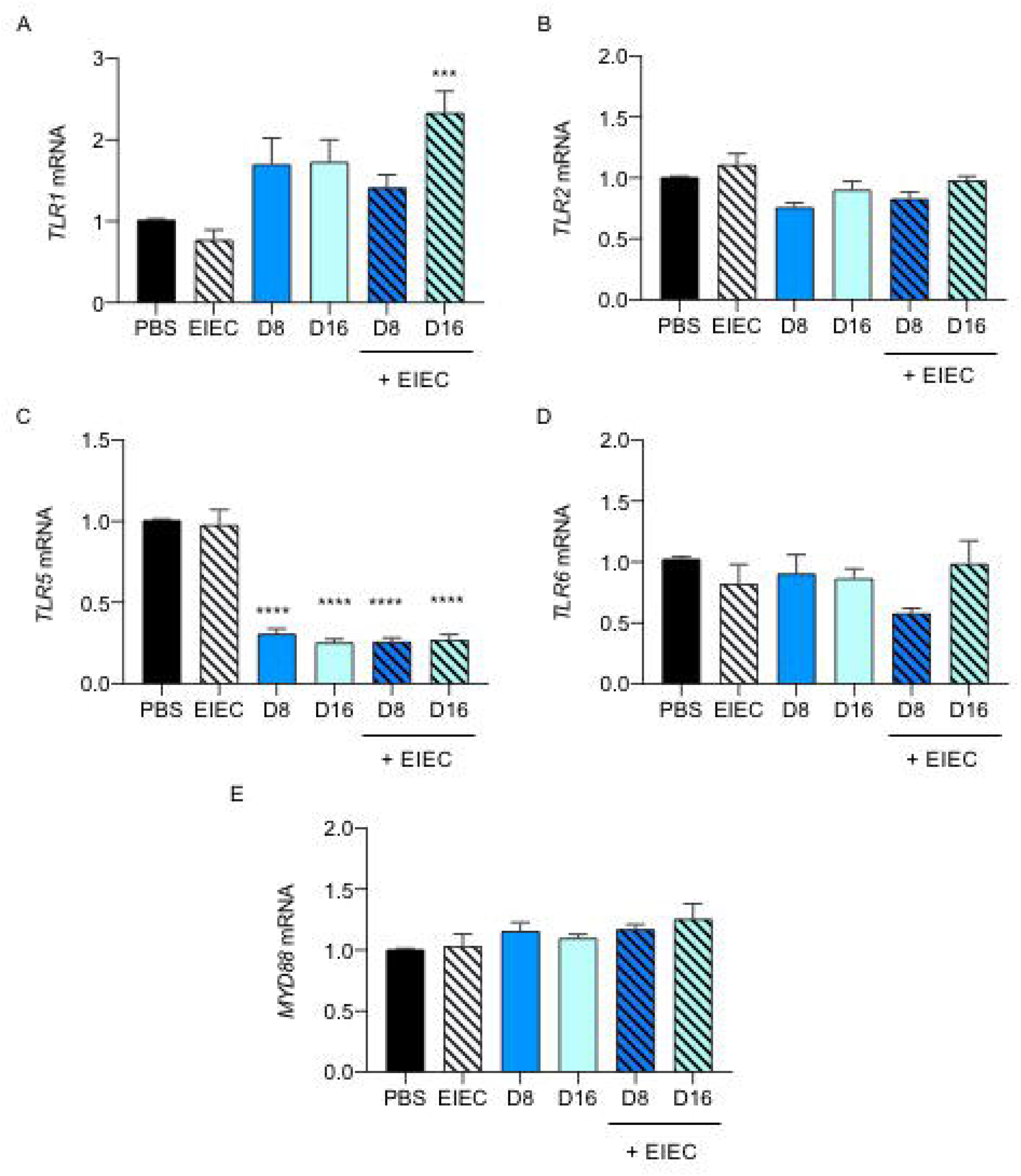
Effects of 24 h DON incubation without or with EIEC post-infection (1 h) on toll-like receptor (TLR) and *MYD88* gene expression. (A-E) *TLR1, TLR2, TLR5, TLR6* and *MYD88* mRNA expression was measured by qPCR, with *GAPDH* as the internal control. Results were shown as mean of ± SEM from five independent experiments. *, ** *p*<0.05, 0.01 compared to PBS control. One-way ANOVA post Dunn’s test.

### 3.6 DON and EIEC tended to activate the NF-κB p65 signalling pathway

p65 (RELA) and p50 (NF-κB1) is the most commonly found heterodimer complex of NF-κB, which participate in nuclear translocation and activation of NF-κB to regulate gene expression and thus major cellular functions (Garg et al. 2013). Post-translational modifications of NF-κB, especially of the RELA subunit, further enhance the NF-κB function as a transcription factor (Huang et al. 2010). Therefore, in our study, qPCR and western blot analyses were used to quantify *RELA* (or NF-κB p65) levels as an indicator for NF-κB activation in response to DON and EIEC contamination (Fig. 7). Treatment with DON alone resulted in an ascending trend in *RELA* mRNA expression. Addition of EIEC further increased the mRNA expression (*p*<0.01) (Fig. 7A). Results obtained from western blot also revealed similar patterns for NF-κB protein expression. Up-regulation of NF-κB p65 protein expression was observed in cells infected with EIEC without and with DON pre-treatment (Fig. 7B).

**Fig. 7.**
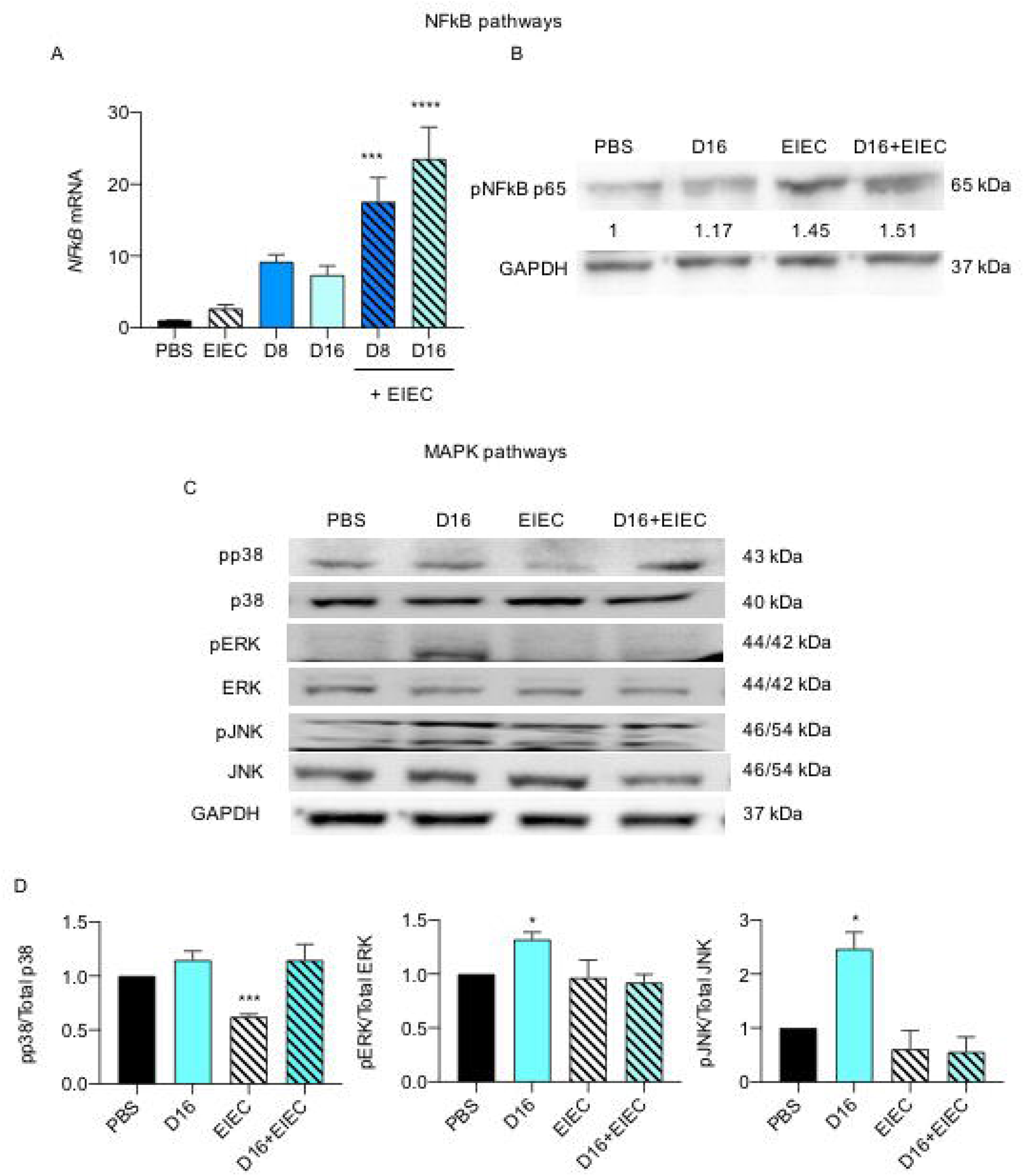
Effects of 24 h DON incubation without or with EIEC post-infection (1 h) on NF-κB and MAPK signalling pathways. (A) *RELA* mRNA expression was measured by qPCR, with *GAPDH* as the internal control. Results were shown as mean of ± SEM from six independent experiments. (B) NF-κB p65 protein expression was measured by Western blotting, with GAPDH as the internal control. Representative photos of western blotting of NF-κB p65 and GAPDH. Quantification of Western blot compared to PBS control from three independent experiments. Inset shows group means. (C) Protein samples were also analysed by Western blot with phospho-p38, JNK and ERK antibodies. The total MAPK levels were used as an internal control. Representative photos of western blotting of MAPKs and GAPDH. Results were shown as mean of ± SEM from four independent experiments. *, ***, **** *p*<0.05, 0.001 and 0.0001 compared to PBS control. One-way ANOVA post Dunn’s test.

### 3.7 EIEC inhibited DON-induced MAPK signalling pathway activation

Previous studies have shown the involvement of MAPK signalling pathways in DON-induced inflammation in intestinal cells (Van De Walle et al. 2008, Van De Walle et al. 2010). In this study the three classical MAPK signalling pathways, JNK, p38 MAPK and extracellular signal-regulated kinase (ERK) were investigated in DON treated cells with and without EIEC post-infection (Fig. 7C). Western blot data showed that DON treatment alone induced significantly the phosphorylation of ERK and JNK, but not p38. However, addition of EIEC to DON treated cells inhibited these MAPK signalling pathways activation. EIEC alone also significantly inhibited p38 signalling pathways but no change in p38 phosphorylation protein level was observed in cells pre-treated with DON.

## 4. Discussion

The present study was the first to investigate the effects of low and relevant concentrations of DON on intestinal susceptibility to acute (1-2 h) EIEC infection. The effects of DON and EIEC contamination on mucin, cytokines and related signal transduction pathways were examined in the intestinal epithelial cells as part of the local immune system. The concentrations of DON are in accordance with the levels probably encountered in the gastrointestinal tract of animals or human tissues after consumption of food or feed contaminated with DON (Sergent et al. 2006). Assuming that DON ingested in one meal is diluted in 1 litre of gastrointestinal fluid and is totally bio-accessible, the *in vitro* concentrations to be used in this study correspond to food contamination ranging from 1.18 mg/kg to 4.72 mg/kg of DON (Van De Walle et al. 2008). The infection protocol (MOI and EIEC treatment duration) was established based on a preliminary experiment in our laboratory. Similar infection protocols were also adopted by other investigators (Resta-Lenert et al. 2003, Ganan et al. 2010, Resta-Lenert et al. 2011).

Bacterial adherence to host cells is the initial crucial step towards colonization and establishment of infection within the host (Torres et al. 2003). In the present study, DON increased the adhesion of EIEC on intestinal epithelial cells (IECs) but caused a reduced invasion into IECs. This may be attributed to the induction of mucin gene and protein expression in IECs as demonstrated in our study. Indeed, numerous studies have shown altered mucin expression in chronic intestinal inflammatory diseases and cancer, both in animal models and patient cohorts (Ho et al. 1993, Reis et al. 1999, Ho et al. 2006, Longman et al. 2006, Moehle et al. 2006, Heazlewood et al. 2008, Larsson et al. 2011). Both secretory and membrane-bound mucins are important constituents of the physicochemical barrier for the protection of the epithelial cell surface against undesirable harmful pathogens (Liévin-Le Moal et al. 2006). Over-expression and hyper-secretion of the secretory mucins, in particular, MUC5AC and MUC5B are two of the important characteristics of the inflammatory process in mucosa. Previous studies conducted by our laboratory indicate the modulation of biosynthesis of MUC5AC and MUC5B following exposure to DON in differentiated Caco-2 cells (Wan et al. 2014). However, no data are available concerning the effects of DON and EIEC on mucin production. Our finding demonstrated that MUC5AC and MUC5B mRNA was significantly increased upon DON and EIEC treatment. The rapid elevation of secretory mucin in responses to xenobiotics and bacterial infection is crucial for protecting intestine against pathogens and its metabolites (Snyder et al. 1987).

DON is known for its ability to activate signalling pathway and gene expression in goblet cells. Several studies indicated the potential involvement of mitogen-activated protein kinases (MAPKs). The initial binding of DON to ribosome leads to the activation of protein kinase R (PKR) that in turn causes the activation of the MAPKs and subsequently up-regulates human *MUC5AC* gene transcription. Less is known about the regulation of MUC5B expression. However, based on the possible common regulatory mechanism between MUC5AC and MUC5B (Moniaux et al. 2001), it is possible that *MUC5B* mRNA expression are regulated by MAPKs activation as well.

There is increasing evidence for the role of membrane-bound mucins in maintaining intestinal mucosal integrity. Among all identified membrane-bound mucins, MUC3 and MUC17 are the membrane-bound mucins that are moderately expressed in the colon (Hattrup et al. 2008) and abundantly in both goblet cells and enterocytes of the small intestine (Ho et al. 1993, Kim et al. 2010). On the other hand, MUC1 and MUC4 are also expressed in normal intestinal tissues, but they are markedly upregulated in response to bacterial infection (McAuley et al. 2007, Lindén et al. 2008). In this study, we have shown a significant induction in *MUC1*, *MUC4* and *MUC17* but not *MUC3* mRNA in cells with DON and EIEC co-exposure. This was in agreement with a previous study that also demonstrated the protective role of MUC17 in protection of the intestinal mucosa against an EIEC strain (Resta-Lenert et al. 2011). MUC17 contributes significantly to maintaining cell homeostasis and modulating chronic inflammatory responses by activating signalling pathways associated with inflammation and cancer. It was postulated that NF-κB contributes, at least partly to the mucin regulation because all intestinally expressed mucin genes contain a potential or experimentally proven binding site for NF-κB (Moehle et al. 2006). The NF-κB regulatory pathway plays an important role in cell activation and production of diverse inflammatory mediators, including a variety of cytokines and chemokines (Hayden et al. 2004). But of course, NF-κB is not the only transcriptional regulator influencing mucin expression. Further studies are necessary to understand the mechanisms controlling the expression of mucin.

Besides acting as a physical barrier, IECs are able to express and produce important mediators of inflammation such as cytokines and chemokines and other signal molecules like TLRs that are important for host defence and bacterial recognition (e.g. lipopolysaccharides (LPS) from gram-negative bacteria) (Arce et al. 2010). TLRs are expressed not only on immune cells but also non-immune cells such as epithelial cells. They can be rapidly induced in response to pathogens, cytokines and environmental stresses. TLR-1, -2, -4, -5 and -6 are expressed on the cell surface, which are implicated in the recognition of microbial membrane components for antimicrobial host defence (Akira et al. 2006). In this study, we showed that upon the treatment with DON and EIEC, the mRNA expression of TLRs was differentially modulated. DON without or with EIEC post-treatment induced *TLR1* gene expression but suppressed *TLR5* expression. *TLR2* and *TLR6*, however showed no significant up-regulation in our present study. This indicates that TLR-1, but not TLR-2 and -6 signalling, was involved in the induction of the early inflammatory responses by EIEC/DON in Caco-2 cells. In contrast, the downregulation of *TLR5* mRNA may function to attenuate excessive inflammatory responses due to DON and EIEC co-exposure. Surprisingly, very low or no expression of TLR-4 was present in any of treatments in Caco-2, and thus respond minimally to EIEC or DON. This is in contrast to other studies which showed induction of TLR expression in response to bacteria toxin (LPS from *Salmonella typhimurium*) in swine intestinal epithelial cell lines (IPEC-J2 and IPI-2I) (Arce et al. 2010), in bovine intestinal epithelial cells following *E. coli* 987P infection (Takanashi et al. 2013), as well as in IPEC-1 cells following enterotoxigenic *E. coli* (ETEC-O149) strain K88 treatment (Taranu et al. 2015). The decrease of TLR-4 and other TLRs (-2, -3, -6) were observed in porcine epithelial cells, macrophages, mesenteric lymph nodes and spleen of mice under the effects of mycotoxins (e.g., T-2 toxins, DON and ZEA) (Seeboth et al. 2012, Islam et al. 2013, Taranu et al. 2015). All these results indicate that differential regulation of TLR gene expression may contribute to inflammatory immune response against bacterial infection in intestinal epithelial cells. However, EIEC treatment for 1 h after DON treatment did not result in more changes in most of the TLRs expression in the present study, implying DON is the major contributing factor for the immune responses.

MyD88 is one of the most important adaptor molecules for inflammatory signalling pathways. MyD88 mediates the activation of TLRs and IL-1R and leads to the production of proinflammatory cytokines through the activation of NF-κB and MAPK signalling pathways (Akira et al. 2006). Here we showed that *RELA* but not *MYD88* mRNA was significantly increased in cells after DON with or without post-challenge with EIEC, in comparison to the unchallenged cells, indicating that NF-κB instead of MyD88 played a more important role in regulating the inflammatory responses induced by DON and EIEC. NF-κB transcription factor plays a critical role in regulation of immune, inflammatory and acute phase responses and is also implicated in the control of cell proliferation and programmed cell death (Aggarwal et al. 2004). NF-κB is harmful to the host when excessively or improperly activated. The ability of DON to influence NF-κB activation has been extensively reported (Van De Walle et al. 2008, Krishnaswamy et al. 2010, Kalaiselvi et al. 2013, Del Regno et al. 2015, Adesso et al. 2017). In this study, we reported that DON increased NF-κB activation during inflammation. EIEC treatment for 1 h after DON treatment caused a higher up-regulation of *RELA* mRNA and NF-κB protein. It is evident that activation of NF-κB is followed by a series of events, leading to the activation of signalling pathways, including MAPKs that are crucial for regulating inflammation and producing inflammatory factors (Van De Walle et al. 2008, Van De Walle et al. 2010). Activation of these signalling cascades could lead to the production of pro-inflammatory cytokines. In the present work, we found that EIEC alone induced the mRNA expression of *IL8* and *TNFA*. Derangement of cytokine production by bacterial infection can lead to chronic inflammatory conditions (Karin et al. 2006). However, it is surprising to show that DON treatment significantly downregulated the mRNA expression of *IL1B IL8* and *TNFA*. This result is in agreement with a previous report by Ghareeb *et al*., which found that in broiler chickens, chronic administration of DON for 5 weeks resulted in significant down-regulation of certain cytokines, such as *IFNG* and *IL1B* mRNA in jejunal tissues (Ghareeb et al. 2013). Similar suppression of splenic *IFNG* and *IL1B* mRNA was also observed in another study in pigs following DON exposure (Cheng et al. 2006). DON is known to be either suppress or stimulate immunological responses, depending on the dose, time and duration of exposure (Ghareeb et al. 2013). In this context, it becomes evident that DON has a modulating effect on the innate immune response. DON could modify the gene expression of cytokines, and thus may affect the susceptibility of human and animals to disease. In spite of the lack of quantifying the levels of proteins that are actually translated from the observed mRNA transcripts, this study is the first to present significant modulation of different pro-inflammatory cytokine mRNA expression in IECs and this might merit further investigation of the mechanisms in relation to the functional relevance of mRNA expression such as by determining the protein levels of the selected pro-inflammatory cytokines by utilizing the quantitative sandwich enzyme immunoassay (ELISA).

Moreover, to determine whether MAPK signalling pathways were involved in the immune responses in cells upon DON and EIEC treatment, the three MAPKs (JNK, p38 MAPK and ERK) were investigated. Consistent with other previous studies, DON induced phosphorylation of JNK and ERK proteins. Addition of EIEC to DON-pre-treated cells, however, suppressed DON-induced phosphorylation of JNK and ERK. DON alone did not induce p38 MAPK phosphorylation but EIEC alone inhibited the p38 MAPK signalling pathway. Although it is evident that MAPK plays an important role in immune responses to *E. coli* infection (Wang et al. 2007, Zhuang et al. 2017), its role in the adherence and internalization of bacteria into epithelial cells was unclear. It is postulated that such deactivation of MAPK pathways may counteract the adhesion and invasion of bacteria into the cells, which are the major contributing factors to intestinal infection and inflammation (Liu et al. 2012).

In conclusion, the above observations provide a context for the present study, suggesting that exposure to DON could be a predisposing factor to infectious disease. IECs are able to generate a rapid immune response against DON and EIEC contaminants when they act alone or in combination. The results also suggested the potential involvement in secretory MUC5AC mucins and membrane bound MUC4 and MUC17 mucins in modifying the attachment and invasion of EIEC and thus affecting the susceptibility to EIEC infection. IECs are able to express and produce important mediators of inflammation such as cytokines and other signal molecules like TLRs that are important for host defence and bacterial recognition. The augmented mucin production and inflammatory stimulation might be a consequence of activation of NF-κB signalling pathway. DON exposure also activated the MAPK signalling molecules, including ERK and JNK through phosphorylation. However, addition of EIEC to DON pre-treated cells inhibited MAPK signalling pathway which might help protecting intestinal epithelial cells from further damages caused by bacterial infection. A summary of the mechanisms of host defence responses against DON and EIEC co-exposure was depicted in Fig. 8. Nevertheless, further studies are necessary to examine different bacterial infection scenarios and to identify the complex mechanism(s) by which this mycotoxin acts on the intestinal tract to modulate invasion and colonization by opportunistic pathogens by using molecular approaches, such as high-throughput mRNA sequencing and proteomics. Epidemiological studies are also needed to assess the extent to which DON are involved in the development of infectious diseases in humans.

**Fig. 8.**
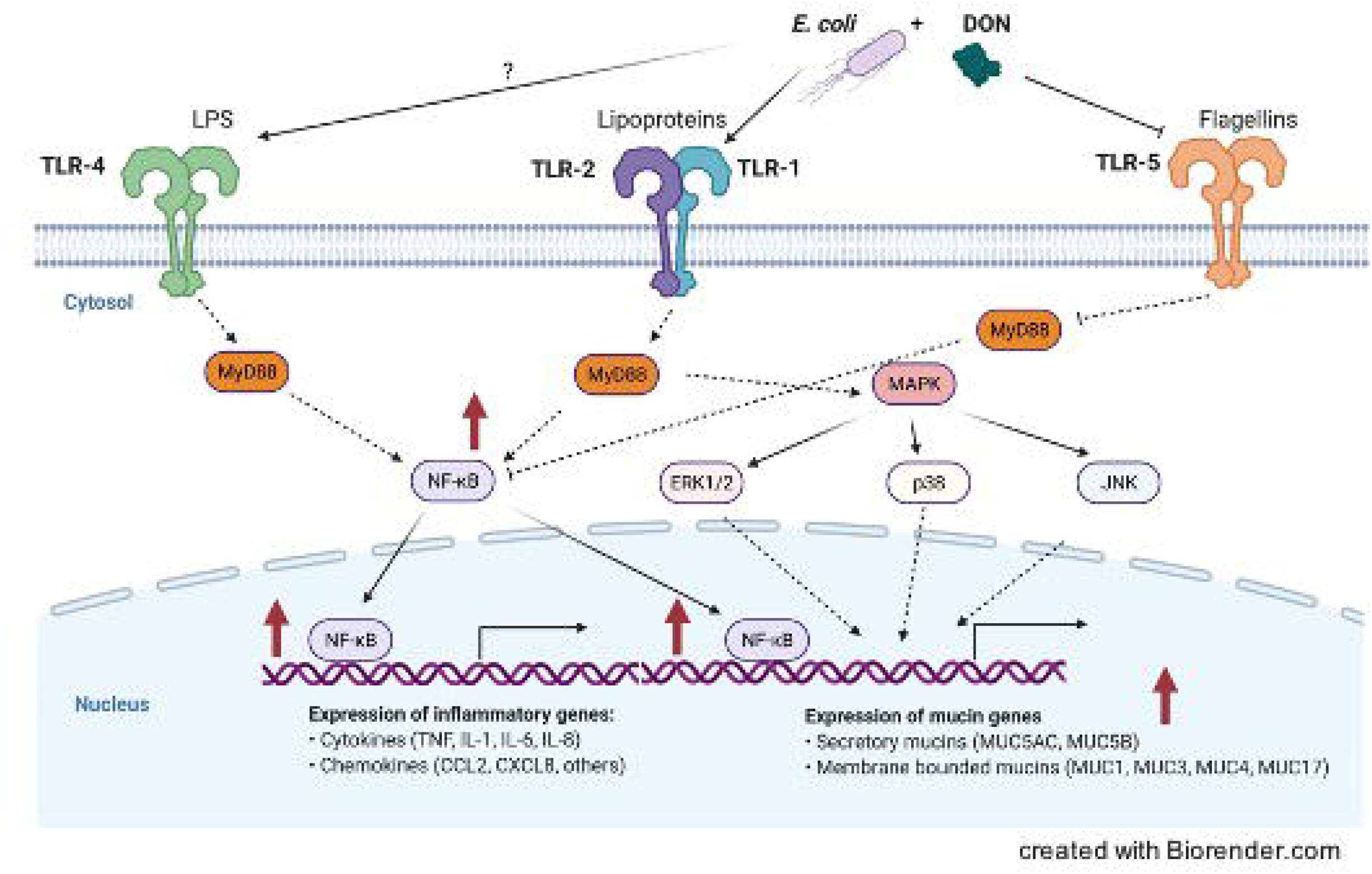
A summary of the proposed mechanisms of host defence responses against DON and EIEC co-exposures. Upon the exposure to DON and EIEC, intestinal epithelial cells are able to express and produce important mediators of inflammation such as cytokines and modulate other signal molecules like TLRs that are important for host defence and bacterial recognition. The augmented mucin production and inflammatory stimulation might be a consequence of activation of NF-κB signalling pathway. Infection of DON pre-treated cells with EIEC inhibited MAPK signalling pathways which might help protecting intestinal epithelial cells from further damages caused by bacterial infection.

## References

Adesso, S., A. Quaroni, A. Popolo, L. Severino and S. Marzocco (2017). “The food contaminants nivalenol and deoxynivalenol induce inflammation in intestinal epithelial cells by regulating reactive oxygen species release.” Nutrients 9(12): 1343.

Aggarwal, B. B., Y. Takada, S. Shishodia, A. M. Gutierrez, O. V. Oommen, H. Ichikawa, Y. Baba and A. Kumar (2004). “Nuclear transcription factor NF-kappa B: role in biology and medicine.” Indian J Exp Biol.

Akira, S., S. Uematsu and O. Takeuchi (2006). “Pathogen recognition and innate immunity.” Cell 124(4): 783–801.

Antonissen, G., F. Van Immerseel, F. Pasmans, R. Ducatelle, G. P. J. Janssens, S. De Baere, K. C. Mountzouris, S. Su, E. A. Wong, B. De Meulenaer, M. Verlinden, M. Devreese, F. Haesebrouck, B. Novak, I. Dohnal, A. Martel and S. Croubels (2015). “Mycotoxins deoxynivalenol and fumonisins alter the extrinsic component of intestinal barrier in broiler chickens.” J Agric Food Chem 63(50): 10846–10855.

Arce, C., M. Ramírez-Boo, C. Lucena and J. Garrido (2010). “Innate immune activation of swine intestinal epithelial cell lines (IPEC-J2 and IPI-2I) in response to LPS from *Salmonella typhimurium*.” Comp Immunol Microbiol Infect Dis 33: 161–174.

Boudeau, J., A.-L. Glasser, E. Masseret, B. Joly and A. Darfeuille-Michaud (1999). “Invasive ability of an *Escherichia coli* strain isolated from the ileal mucosa of a patient with Crohn’s disease.” Infect Immun 67(9): 4499–4509.

Cheng, Y.-H., C.-F. Weng, B.-J. Chen and M.-H. Chang (2006). “Toxicity of different *Fusarium* mycotoxins on growth performance, immune responses and efficacy of a mycotoxin degrading enzyme in pigs.” Anim Res 55(6): 579–590.

Croxen, M. A., R. J. Law, R. Scholz, K. M. Keeney, M. Wlodarska and B. B. Finlay (2013). “Recent advances in understanding enteric pathogenic *Escherichia coli*.” Clin Microbiol Rev 26(4): 822–880.

Dąbek, J., A. Kułach and Z. Gąsior (2010). “Nuclear factor kappa-light-chain-enhancer of activated B cells (NF-κB): a new potential therapeutic target in atherosclerosis?” Pharmacol Rep 62(5): 778–783.

Del Regno, M., S. Adesso, A. Popolo, A. Quaroni, G. Autore, L. Severino and S. Marzocco (2015). “Nivalenol induces oxidative stress and increases deoxynivalenol pro-oxidant effect in intestinal epithelial cells.” Toxicol Appl Pharm 285(2): 118–127.

Eaves-Pyles, T., C. A. Allen, J. Taormina, A. Swidsinski, C. B. Tutt, G. Eric Jezek, M. Islas-Islas and A. G. Torres (2008). “*Escherichia coli* isolated from a Crohn’s disease patient adheres, invades, and induces inflammatory responses in polarized intestinal epithelial cells.” Int J Med Microbiol 298(5): 397–409.

Fukata, T., K. Sasai, E. Baba and A. Arakawa (1996). “Effect of ochratoxin A on *Salmonella typhimurium*-challenged layer chickens.” Avian Dis: 924–926.

Ganan, M., M. Collins, R. Rastall, A. Hotchkiss, H. Chau, A. Carrascosa and A. Martinez-Rodriguez (2010). “Inhibition by pectic oligosaccharides of the invasion of undifferentiated and differentiated Caco-2 cells by *Campylobacter jejuni*.” Int J Food Microbiol 137(2-3): 181–185.

Garg, M., J. A. Potter and V. M. Abrahams (2013). “Identification of microRNAs that regulate TLR2-mediated trophoblast apoptosis and inhibition of IL-6 mRNA.” PLOS One 8(10): e77249.

Ghareeb, K., W. A. Awad, C. Soodoi, S. Sasgary, A. Strasser and J. Böhm (2013). “Effects of feed contaminant deoxynivalenol on plasma cytokines and mRNA expression of immune genes in the intestine of broiler chickens.” PLOS One 8(8): e71492.

Hattrup, C. L. and S. J. Gendler (2008). “Structure and function of the cell surface (tethered) mucins.” Annu Rev Physiol 70: 431–457.

Hayden, M. S. and S. Ghosh (2004). “Signaling to NF-κB.” Genes Dev 18(18): 2195–2224.

Heazlewood, C. K., M. C. Cook, R. Eri, G. R. Price, S. B. Tauro, D. Taupin, D. J. Thornton, C. W. Png, T. L. Crockford, R. J. Cornall, R. Adams, M. Kato, K. A. Nelms, N. A. Hong, T. H. J. Florin, C. C. Goodnow and M. A. McGuckin (2008). “Aberrant mucin assembly in mice causes endoplasmic reticulum stress and spontaneous inflammation resembling ulcerative colitis.” PLOS Med 5(3): e54.

Ho, S. B., L. A. Dvorak, R. E. Moor, A. C. Jacobson, M. R. Frey, J. Corredor, D. B. Polk and L. L. Shekels (2006). “Cysteine-rich domains of muc3 intestinal mucin promote cell migration, inhibit apoptosis, and accelerate wound healing.” Gastroenterology 131(5): 1501–1517.

Ho, S. B., G. A. Niehans, C. Lyftogt, P. S. Yan, D. L. Cherwitz, E. T. Gum, R. Dahiya and Y. S. Kim (1993). “Heterogeneity of mucin gene expression in normal and neoplastic tissues.” Cancer Res 53(3): 641–651.

Huang, B., X.-D. Yang, A. Lamb and L.-F. Chen (2010). “Posttranslational modifications of NF-kappaB: another layer of regulation for NF-kappaB signaling pathway.” Cell Signal 22(9): 1282–1290.

Ishiyama, M., Y. Miyazono, K. Sasamoto, Y. Ohkura and K. Ueno (1997). “A highly water-soluble disulfonated tetrazolium salt as a chromogenic indicator for NADH as well as cell viability.” Talanta 44(7): 1299–1305.

Islam, M. R., Y. S. Roh, J. Kim, C. W. Lim and B. Kim (2013). “Differential immune modulation by deoxynivalenol (vomitoxin) in mice.” Toxicol Lett 221(2): 152–163.

Jung, H., L. Eckmann, S. Yang, A. Panja, J. Fierer, E. Morzycka-Wroblewska and M. Kagnoff (1995). “A distinct array of proinflammatory cytokines is expressed in human colon epithelial cells in response to bacterial invasion.” J Clin Invest 95: 55–65.

Kalaiselvi, P., K. Rajashree, L. B. Priya and V. V. Padma (2013). “Cytoprotective effect of epigallocatechin-3-gallate against deoxynivalenol-induced toxicity through anti-oxidative and anti-inflammatory mechanisms in HT-29 cells.” Food Chem Toxicol 56: 110–118.

Karin, M., T. Lawrence and V. Nizet (2006). “Innate immunity gone awry: linking microbial infections to chronic inflammation and cancer.” Cell 124(4): 823–835.

Khodaii, Z., S. M. H. Ghaderian and M. M. Natanzi (2017). “Probiotic bacteria and their supernatants protect enterocyte cell lines from enteroinvasive *Escherichia coli* (EIEC) invasion.” Int J Mol Cell Med 6(3): 183.

Kim, Y. S. and S. B. Ho (2010). “Intestinal goblet cells and mucins in health and disease: recent insights and progress.” Curr Gastroenterol Rep 12(5): 319–330.

Krishnaswamy, R., S. N. Devaraj and V. V. Padma (2010). “Lutein protects HT-29 cells against deoxynivalenol-induced oxidative stress and apoptosis: prevention of NF-κB nuclear localization and down regulation of NF-κB and cyclo-oxygenase-2 expression.” Free Radic Biol Med 49(1): 50–60.

Larsson, J. M. H., H. Karlsson, J. G. Crespo, M. E. Johansson, L. Eklund, H. Sjövall and G. C. Hansson (2011). “Altered O-glycosylation profile of MUC2 mucin occurs in active ulcerative colitis and is associated with increased inflammation.” Inflamm Bowel Dis 17(11): 2299–2307.

Liévin-Le Moal, V. and A. L. Servin (2006). “The front line of enteric host defense against unwelcome intrusion of harmful microorganisms: mucins, antimicrobial peptides, and microbiota.” Clin Microbiol Rev 19(2): 315–337.

Lindén, S. K., T. H. J. Florin and M. A. McGuckin (2008). “Mucin dynamics in intestinal bacterial infection.” PLOS One 3(12): e3952.

Liu, Z., Y. Ma, M. P. Moyer, P. Zhang, C. Shi and H. Qin (2012). “Involvement of the mannose receptor and p38 mitogen-activated protein kinase signaling pathway of the microdomain of the integral membrane protein after enteropathogenic *Escherichia coli* infection.” Infect Immun 80(4): 1343–1350.

Livak, K. J. and T. D. Schmittgen (2001). “Analysis of relative gene expression data using real-time quantitative PCR and the 2−ΔΔCT method.” Methods 25(4): 402–408.

Longman, R. J., R. Poulsom, A. P. Corfield, B. F. Warren, N. A. Wright and M. G. Thomas (2006). “Alterations in the composition of the supramucosal defense barrier in relation to disease severity of ulcerative colitis.” J Histochem Cytochem 54(12): 1335–1348.

Maciorowski, K. G., P. Herrera, F. T. Jones, S. D. Pillai and S. C. Ricke (2007). “Effects on poultry and livestock of feed contamination with bacteria and fungi.” Anim Feed Sci Tech 133(1): 109–136.

McAuley, J. L., S. K. Linden, C. W. Png, R. M. King, H. L. Pennington, S. J. Gendler, T. H. Florin, G. R. Hill, V. Korolik and M. A. McGuckin (2007). “MUC1 cell surface mucin is a critical element of the mucosal barrier to infection.” J Clin Invest 117(8): 2313–2324.

Moehle, C., N. Ackermann, T. Langmann, C. Aslanidis, A. Kel, O. Kel-Margoulis, A. Schmitz-Madry, A. Zahn, W. Stremmel and G. Schmitz (2006). “Aberrant intestinal expression and allelic variants of mucin genes associated with inflammatory bowel disease.” J Mol Med 84(12): 1055–1066.

Moniaux, N., F. Escande, N. Porchet, J. Aubert and S. Batra (2001). “Structural organization and classification of the human mucin genes.” Front Biosci 6: D1192–1206.

Moon, Y. and J. Pestka (2002). “Vomitoxin-induced cyclooxygenase-2 gene expression in macrophages mediated by activation of ERK and p38 but not JNK mitogen-activated protein kinases.” Toxicol Sci 69: 373–382.

Moon, Y. and J. Pestka (2003). “Cyclooxygenase-2 mediates interleukin-6 upregulation by vomitoxin (deoxynivalenol) *in vitro* and *in vivo*.” Toxicol Appl Pharm 187: 80–88.

Nolan, T., R. E. Hands and S. A. Bustin (2006). “Quantification of mRNA using real-time RT-PCR.” Nat Protocols 1(3): 1559–1582.

Oswald, I., C. Desautels, J. Laffitte, S. Fournout, S. Peres, M. Odin, P. Le Bars, J. Le Bars and J. Fairbrother (2003). “Mycotoxin fumonisin B1 increases intestinal colonization by pathogenic *Escherichia coli* in pigs.” Appl Environ Microbiol 69: 5870–5874.

Pestka, J., H. Zhou, Y. Moon and Y. Chung (2004). “Cellular and molecular mechanisms for immune modulation by deoxynivalenol and other trichothecenes: unraveling a paradox.” Toxicol Lett 153: 61–73.

Pinton, P., F. Graziani, A. Pujol, C. Nicoletti, O. Paris, P. Ernouf, E. Di Pasquale, J. Perrier, I. P. Oswald and M. Maresca (2015). “Deoxynivalenol inhibits the expression by goblet cells of intestinal mucins through a PKR and MAP kinase dependent repression of the resistin-like molecule β.” Mol Nutr Food Res 59: 1076–1087.

Reis, C. A., L. David, P. Correa, F. Carneiro, C. de Bolós, E. Garcia, U. Mandel, H. Clausen and M. Sobrinho-Simões (1999). “Intestinal metaplasia of human stomach displays distinct patterns of mucin (MUC1, MUC2, MUC5AC, and MUC6) expression.” Cancer Res 59(5): 1003–1007.

Resta-Lenert, S. and K. E. Barrett (2003). “Live probiotics protect intestinal epithelial cells from the effects of infection with enteroinvasive *Escherichia coli* (EIEC).” Gut 52(7): 988–997.

Resta-Lenert, S., S. Das, S. K. Batra and S. B. Ho (2011). “Muc17 protects intestinal epithelial cells from enteroinvasive *E. coli* infection by promoting epithelial barrier integrity.” Am J Physiol-Gastrointest Liver Physiol 300(6): G1144–G1155.

Schneider, C. A., W. S. Rasband and K. W. Eliceiri (2012). “NIH Image to ImageJ: 25 years of image analysis.” Nat Methods 9: 671–675.

Seeboth, J., R. Solinhac, I. P. Oswald and L. Guzylack-Piriou (2012). “The fungal T-2 toxin alters the activation of primary macrophages induced by TLR-agonists resulting in a decrease of the inflammatory response in the pig.” Vet Res 43(1): 35.

Sergent, T., M. Parys, S. Garsou, L. Pussemier, Y.-J. Schneider and Y. Larondelle (2006). “Deoxynivalenol transport across human intestinal Caco-2 cells and its effects on cellular metabolism at realistic intestinal concentrations.” Toxicol Lett 164(2): 167–176.

Snyder, J. D. and A. Walker (1987). “Structure and function of intestinal mucin: developmental aspects.” Int Arch Allergy Immunol 82(3-4): 351–356.

Sobrova, P., V. Adam, A. Vasatkova, M. Beklova, L. Zeman and R. Kizek (2010). “Deoxynivalenol and its toxicity.” Interdiscip Toxicol 3(3): 94–99.

Stoev, S., D. Goundasheva, T. Mirtcheva and P. Mantle (2000). “Susceptibility to secondary bacterial infections in growing pigs as an early response in ochratoxicosis.” Exp Toxicol Pathol 52(4): 287–296.

Tai, J. H. and J. J. Pestka (1988). “Impaired murine resistance to *Salmonella typhimurium* following oral exposure to the trichothecene T-2 toxin.” Food Chem Toxicol 26(8): 691–698.

Takanashi, N., Y. Tomosada, J. Villena, K. Murata, T. Takahashi, E. Chiba, M. Tohno, T. Shimazu, H. Aso, Y. Suda, S. Ikegami, H. Itoh, Y. Kawai, T. Saito, S. Alvarez and H. Kitazawa (2013) “Advanced application of bovine intestinal epithelial cell line for evaluating regulatory effect of lactobacilli against heat-killed enterotoxigenic *Escherichia coli*-mediated inflammation.” BMC Microbiol 13, 54 DOI: 10.1186/1471-2180-13-54.

Taranu, I., D. E. Marin, G. C. Pistol, M. Motiu and D. Pelinescu (2015). “Induction of pro-inflammatory gene expression by *Escherichia coli* and mycotoxin zearalenone contamination and protection by a *Lactobacillus* mixture in porcine IPEC-1 cells.” Toxicon 97: 53–63.

Torres, A. G. and J. B. Kaper (2003). “Multiple elements controlling adherence of enterohemorrhagic *Escherichia coli* O157: H7 to HeLa cells.” Infect Immun 71(9): 4985–4995.

Underhill, D. M. and A. Ozinsky (2002). “Toll-like receptors: key mediators of microbe detection.” Curr Opin Immunol 14(1): 103–110.

Van De Walle, J., A. During, N. Piront, O. Toussaint, Y.-J. Schneider and Y. Larondelle (2010). “Physio-pathological parameters affect the activation of inflammatory pathways by deoxynivalenol in Caco-2 cells.” Toxicol In Vitro 24(7): 1890–1898.

Van De Walle, J., B. Romier, Y. Larondelle and Y. Schneider (2008). “Influence of deoxynivalenol on NF-κB activation and IL-8 secretion in human intestinal Caco-2 cells.” Toxicol Lett 177: 205–214.

Van De Walle, J., B. Romier, Y. Larondelle and Y.-J. Schneider (2008). “Influence of deoxynivalenol on NF-κB activation and IL-8 secretion in human intestinal Caco-2 cells.” Toxicol Lett 177(3): 205–214.

Vandenbroucke, V., S. Croubels, A. Martel, E. Verbrugghe, J. Goossens, K. Van Deun, F. Boyen, A. Thompson, N. Shearer, P. De Backer, F. Haesebrouck and F. Pasmans (2011). “The mycotoxin deoxynivalenol potentiates intestinal inflammation by *Salmonella Typhimurium* in porcine ileal loops.” PLOS One 6(8): e23871.

Vieira, M. A. M., T. A. T. Gomes, A. J. P. Ferreira, T. Knöbl, A. L. Servin and V. Liévin-Le Moal (2010). “Two atypical enteropathogenic *Escherichia coli* strains induce the production of secreted and membrane-bound mucins to benefit their own growth at the apical surface of human mucin-secreting intestinal HT29-MTX cells.” Infect Immun 78(3): 927–938.

Wan, L.-Y. M., K. J. Allen, P. C. Turner and H. El-Nezami (2014). “Modulation of mucin mRNA (MUC5AC and MUC5B) expression and protein production and secretion in Caco-2/HT29-MTX co-cultures following exposure to individual and combined Fusarium mycotoxins.” Toxicol Sci 139(1): 83–98.

Wan, L.-Y. M., C.-S. J. Woo, P. C. Turner, J. M.-F. Wan and H. El-Nezami (2013). “Individual and combined effects of *Fusarium* toxins on the mRNA expression of pro-inflammatory cytokines in swine jejunal epithelial cells.” Toxicol Lett 220(3): 238–246.

Wang, J. H., Y. J. Zhou, P. He and B. Y. Chen (2007). “Roles of mitogen-activated protein kinase pathways during Escherichia coli-induced apoptosis in U937 cells.” Apoptosis 12(2): 375–385.

Xue, Y., H. Zhang, H. Wang, J. Hu, M. Du and M.-J. Zhu (2014). “Host inflammatory response inhibits *Escherichia coli* O157:H7 adhesion to gut epithelium through augmentation of mucin expression.” Infect Immun 82(5): 1921–1930.

Zhuang, X., Z. Chen, C. He, L. Wang, R. Zhou, D. Yan and B. Ge (2017). “Modulation of host signaling in the inflammatory response by enteropathogenic *Escherichia coli* virulence proteins.” Cell Mol Immunmol 14(3): 237–244.

